# Repository-based plasmid design Cost-optimal Gibson Assembly with repository DNA selection and primer generation

**DOI:** 10.1101/792341

**Authors:** Joshua J. Timmons, Doug Densmore

## Abstract

There was an explosion in the amount of commercially available DNA in sequence repositories over the last decade. The number of such plasmids increased from 12,000 to over 300,000 among three of the largest repositories: iGEM, Addgene, and DNASU. The challenge in biodesign remains how to use these and other repository-based sequences effectively, correctly, and seamlessly. This work describes an approach to plasmid design where a plasmid is specified as a DNA sequence or list of features. The proposed software then finds the most cost-effective combination of synthetic and PCR-prepared repository fragments to build the plasmid via Gibson Assembly. It finds existing DNA sequences in user-specified and public DNA databases: iGEM, Addgene, and DNASU. Such a software application is introduced and characterized against all post-2005 iGEM composite parts and all Addgene vectors submitted in 2018 and found to reduce costs by 34% versus a purely synthetic plasmid design approach. The described software will improve current plasmid assembly workflows by shortening design times, improving build quality, and reducing costs.

## INTRODUCTION

Plasmids are a common design element in molecular biology. They are used for studying genes (1), encoding programs (2), and editing human cells (3). Most are simple relative to their bacterial progenitors, containing just an origin of replication, resistance marker – combined together in a “backbone” – and an insert region of exogenous DNA (4). But the exogenous region of plasmids are increasingly complex; genetic circuits in synthetic biology, for example, can comprise more than a dozen separate elements packaged together (2,5,6).

Electronically available plasmid repositories are centers for plasmid sharing (7). They enable cheap, reproducible, and collaborative experimental designs while handling the tedious processes of sample categorization and archiving (8). Plus, they assist researchers required by journals to make their materials publicly available. And, whether by choice or pressure from journals and grant agencies, researchers have been submitting their plasmids to repositories in increasing number. Addgene, for example, grew from zero plasmids in 2004 to more than 70,000 in 2019 and has shipped over a million samples to researchers in 96 countries (8–10). iGEM’s Registry of Standardized Biological Parts increased from 899 to more than 26,000 over the same time period (11,12) (Fig 1). DNASU now has greater than 200,000 plasmids and has shipped over 360,000 clones. Other large repositories include PlasmID (13), of Harvard Medical School, the BCCM/GeneCorner repository of Ghent University in Belgium, and the Fungal Genetics Stock Center (14) of Kansas State University. Plasmid repositories’ inventories will only grow in future years: another 100% increase, like the one that over the last five years, would represent another 300,000 orderable plasmids.

**Fig 1:**
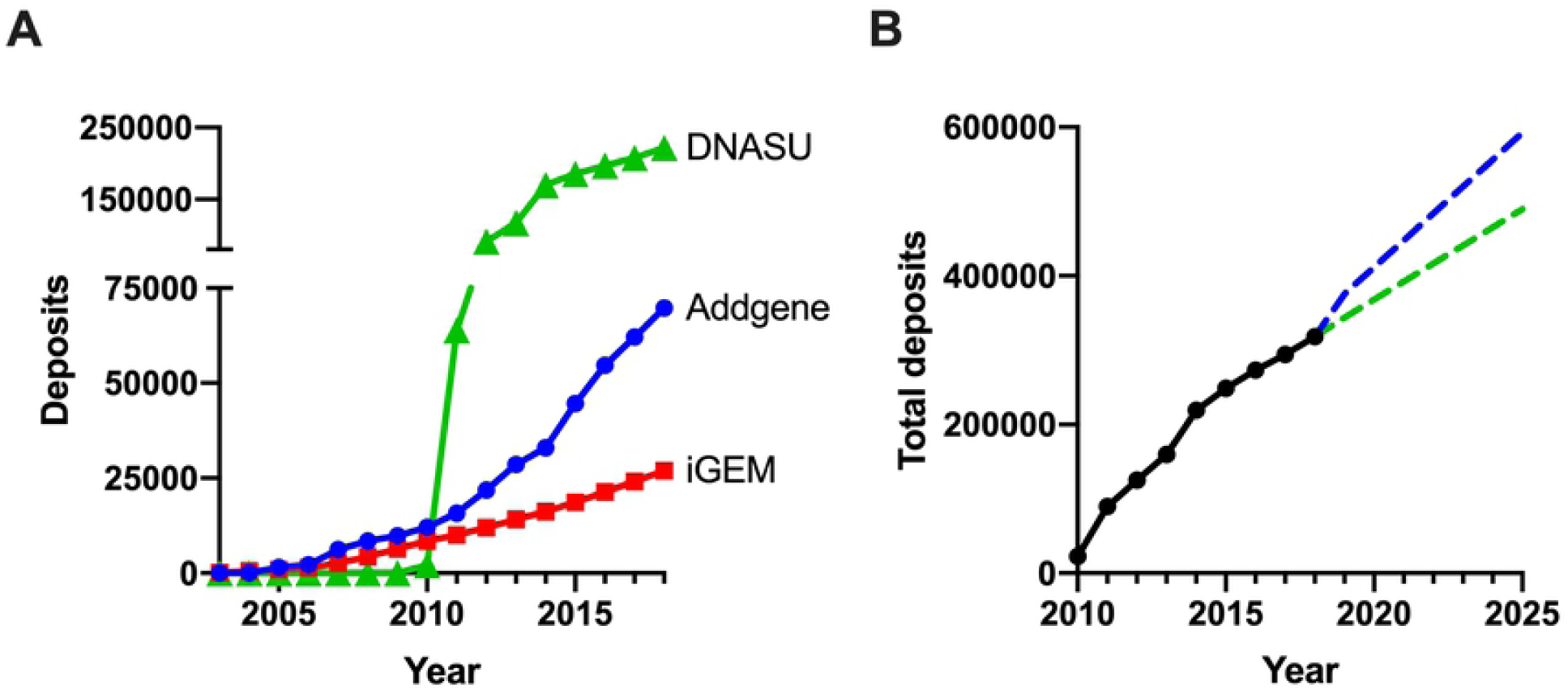
Rapid Growth in DNA Repository Size. (A) Total number of DNA deposits in iGEM, Addgene, and DNASU from 2005 to 2018. (B) Total number of deposits between 2010 and 2018 with linear regression forecasts through 2025 using five (green) or ten (blue) years of past deposit totals. There was a 25-fold increase in the total number of sequence deposits available between 2008 and 2018 and a 99% increase between 2013 and 2018. Continued growth like the last five years would result in roughly 500,000 accumulated deposits between iGEM, Addgene, and DNASU by 2025.

When designing a new plasmid, an experimentalist can assemble it from fragments already in their or their co-workers’ labs (“local” repositories), public DNA repositories, or order it from a synthesis provider (either as a fragment or as a pre-cloned gene). Because combining fragments from local and remote repositories is presently less expensive than synthesis, and because plasmids in repositories may contain the features of biologists’ desired plasmids, we believe that plasmid repositories should be considered in the plasmid design process.

Repurposing DNA in repositories for new plasmids is non-trivial for several reasons. First, sequences in remote repositories are unstandardized – unlike those in the iGEM registry or Golden Gate kits (15–17) – so fragments will require preparation. Second, the researcher must be aware of where the redundant plasmid sequence is stored in remote repositories. Finally, even if a researcher is aware of fragments within a repository that they want to combine, they may fail to find the least expensive combination of repository fragments and synthesized fragments to assemble their plasmid without any off-target primers, replicated fragment junctions, or hairpins in the ends of synthetic fragments (18). For example, the least expensive plasmid design for a user may contain a combination of fragments from Addgene, DNASU, and several short synthetic fragments. Finding and comparing all combinations of fragment sources from multiple repositories and synthesis providers, to build the cheapest plasmid, is beyond the capabilities of a human plasmid designer.

We have designed an application, REPP (an acronym for “repository” and “plasmid”), that addresses the issues raised above. It collates sequences from both public and user repositories and factors in the cost of synthesis to find the cheapest plasmid design possible. REPP circumvents the lack of standardization in the repositories by designing for Gibson Assembly (19–21). It creates homology between adjacent fragments via primers (created with Primer3). Finally, it checks and avoids primers with off-targets, replicated junctions, and synthetic fragments with hairpins in their ends.

There are other tools for preparing a list of DNA fragments for Gibson Assembly. They include, but are not limited to, j5 (22), Benchling (23), SnapGene (24), and Geneious (25). Each simulates an assembly of fragments and outputs a list of fragments with primers for preparation. But each depends on the user knowing beforehand which combination of fragments they want in the plasmid design – despite a less expensive solution possibly existing in the freezer or among the plasmids in external repositories. REPP incorporates user repositories, public repositories, and synthesis – applying costs to each – to build minimum cost solutions that abide by Gibson Assembly design rules.

REPP reverses the design process of the aforementioned plasmid design tools. Rather than choosing fragments and annealing them into a plasmid, the user specifies a plasmid’s sequence or features and REPP decomposes it into a set of fragments and primers. A user specifies their plasmid and REPP finds the fragments and primers to create the plasmid with the least expensive cost. In software engineering terms, REPP has a declarative interface rather than an imperative one (26); users specify the plasmid they want rather than the fragments to put it together. We believe this higher-level approach to plasmid design will be more intuitive and powerful for users today and automated labs of the future where sequences are generated automatically without human intervention. REPP includes fragment-based plasmid specification as well, but it is not discussed here.

A recently released but comparable tool is DNA Weaver (27). Like the tool discussed here, it accepts a “target” sequence as an input and outputs the fragments, existing or newly synthesized, to assemble it. The greatest difference between DNA Weaver and REPP is that DNA Weaver constructs linear sequences while REPP builds circular plasmids. Furthermore, DNA Weaver lacks feature-based plasmid design and the embedded sequence repositories included in REPP but has graphical reports describing build output and has support for Golden Gate Assembly, neither of which are supported by REPP.

REPP is open-source on Github (28) and pre-compiled binaries are available for Linux, MacOS, and Windows from SourceForge (29). Primer3, BLAST, and BLAST databases for iGEM, DNASU, and Addgene are bundled with the application.

## MATERIALS AND METHODS

### Design Overview

Users of REPP specify a target plasmid as either a DNA sequence or as a list of feature names. They also choose a set of fragment databases (BLAST databases) to use as build sources: Addgene, DNASU, and iGEM are included, and local (same machine) BLAST databases are supported as well. REPP then finds the minimum cost design instructions for Gibson Assembly. It outputs the name and repository URL of build fragments used to construct portions of the target plasmid. It creates primers for amplifying the build fragments, with junctions for neighboring fragments, as well as a list of synthetic fragments if necessary. It chooses PCR fragments and synthetic fragments so the designs are the least expensive possible given the costs and constraints in a user specified configuration file. A high-level flow of REPP is detailed in Fig 4. For details about REPP’s implementation, we invite researchers to visit the application’s code repository.

**Fig 2:**
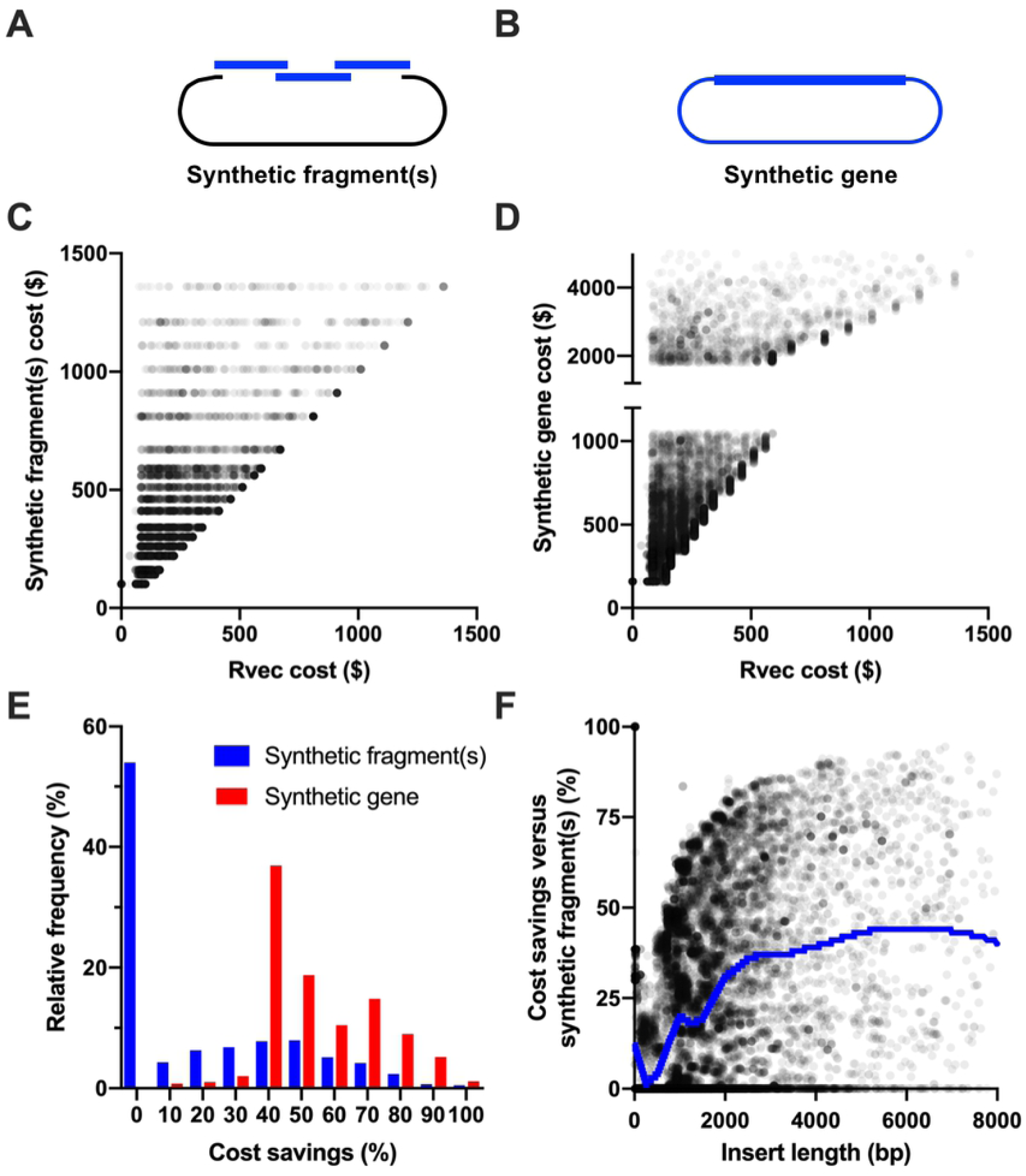
Plasmid Design of iGEM Parts with Preexisting iGEM Parts. REPP’s design of iGEM composite devices with 98% identity, synthesis with IDT, and iGEM itself as the sole fragment source repository. (A) An alternative to REPP’s designs of each plasmid was their assembly with synthetic fragment(s) alone without PCR’ed sub-fragments. If an iGEM part was too long for a single synthetic fragment, it was divided into multiple synthetic sub-fragments. (B) Another alternative build approach was ordering the iGEM part in a “synthetic gene” where the part is delivered in a pre-cloned plasmid. (C) Cost comparison between plasmids designed with synthetic fragment(s) inserted into linearized pSB1A3 and REPP’s plasmid designs for all iGEM plasmids. For the top quartile of the plasmid designs, REPP’s solution was at least 39% less expensive than synthetic fragments alone. (D) Similar cost comparison between REPP’s plasmid designs and synthetic gene costs. REPP’s solution was always cheaper, though often included only synthetic fragments. (E) Frequency distributions for the cost savings of REPP’s plasmid designs versus synthetic fragment(s) (blue) and synthetic genes (red). (F) Length of the iGEM part (insert) versus REPP’s cost savings against synthetic fragment-only plasmid design. A Lowess curve, blue, was created with a windows size of 15%. REPP’s savings were greater than 25% for all plasmid designs greater than 2,000bp and averaged around 37% for plasmids with 3,000bp iGEM parts.

**Fig 3:**
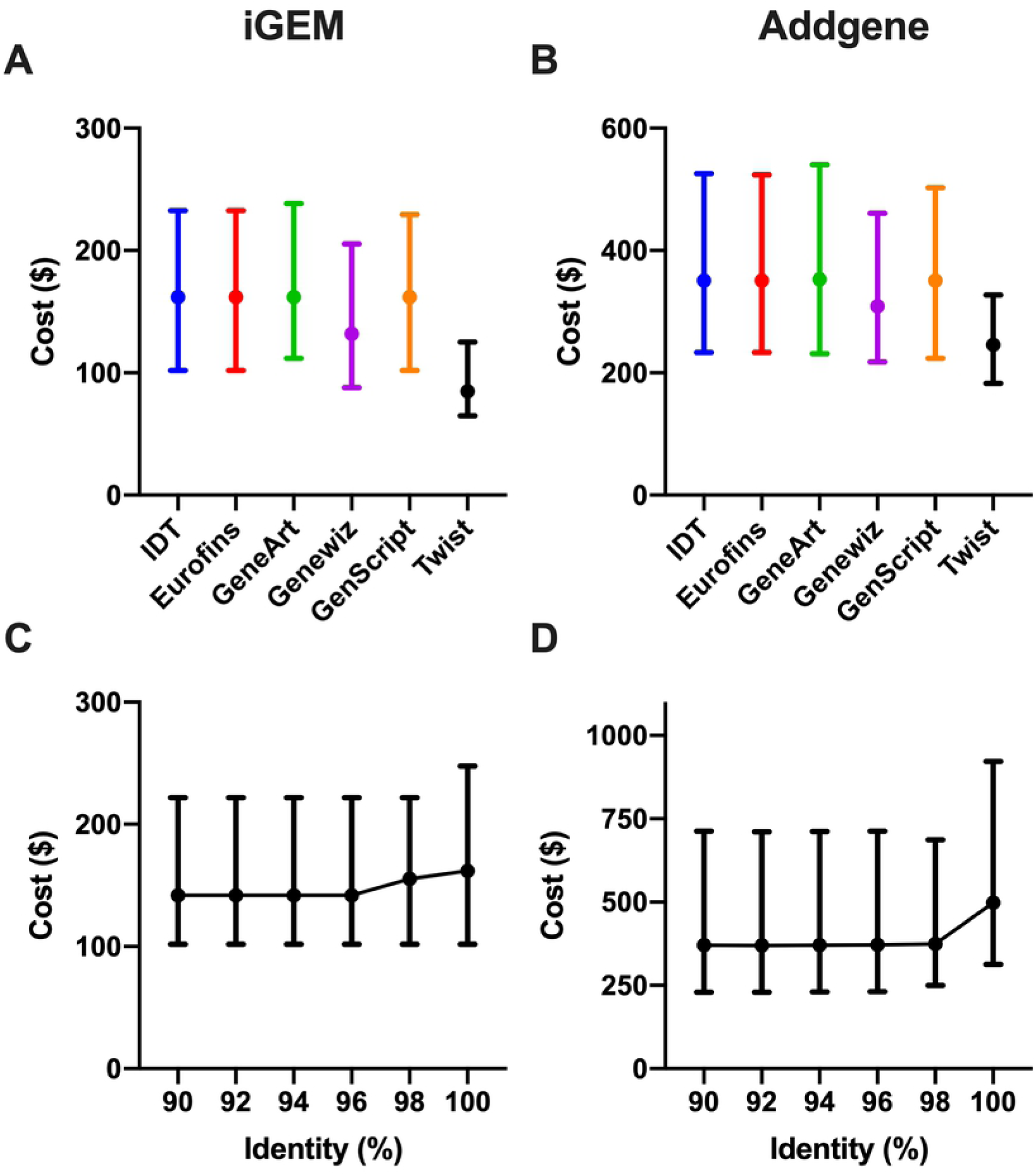
Cost Sensitivities. Plasmid assembly costs for the iGEM and Addgene datasets in response to varied synthesis providers and percentage identities. Median and interquartile range of REPP designed plasmid costs using synthesis costs corresponding to IDT, Eurofins, GeneArt, Genewiz, GenScript, and Twist for the iGEM (A) and Addgene (B) datasets. Synthesis costs were from February 2019. Less expensive synthesis costs like those Twist corresponded with cheaper overall plasmid costs from REPP. REPP’s specification file makes the cost adaptable to future price changes. In (A), the dataset is the same as Fig 2: composite iGEM parts inserted into pSB1A3 digested with PstI and a 98% sequence identity requirement. The plasmids in (B) were all uploaded to the Addgene repository in 2018. Median and interquartile range of plasmid costs in response to varied percentage identities for the iGEM (C) and Addgene 2018 (B) datasets. There was an immediate drop in median plasmid cost at 98% identity for both the iGEM and Addgene datasets which lead to its adoption as the default percentage identity.

**Fig 4:**
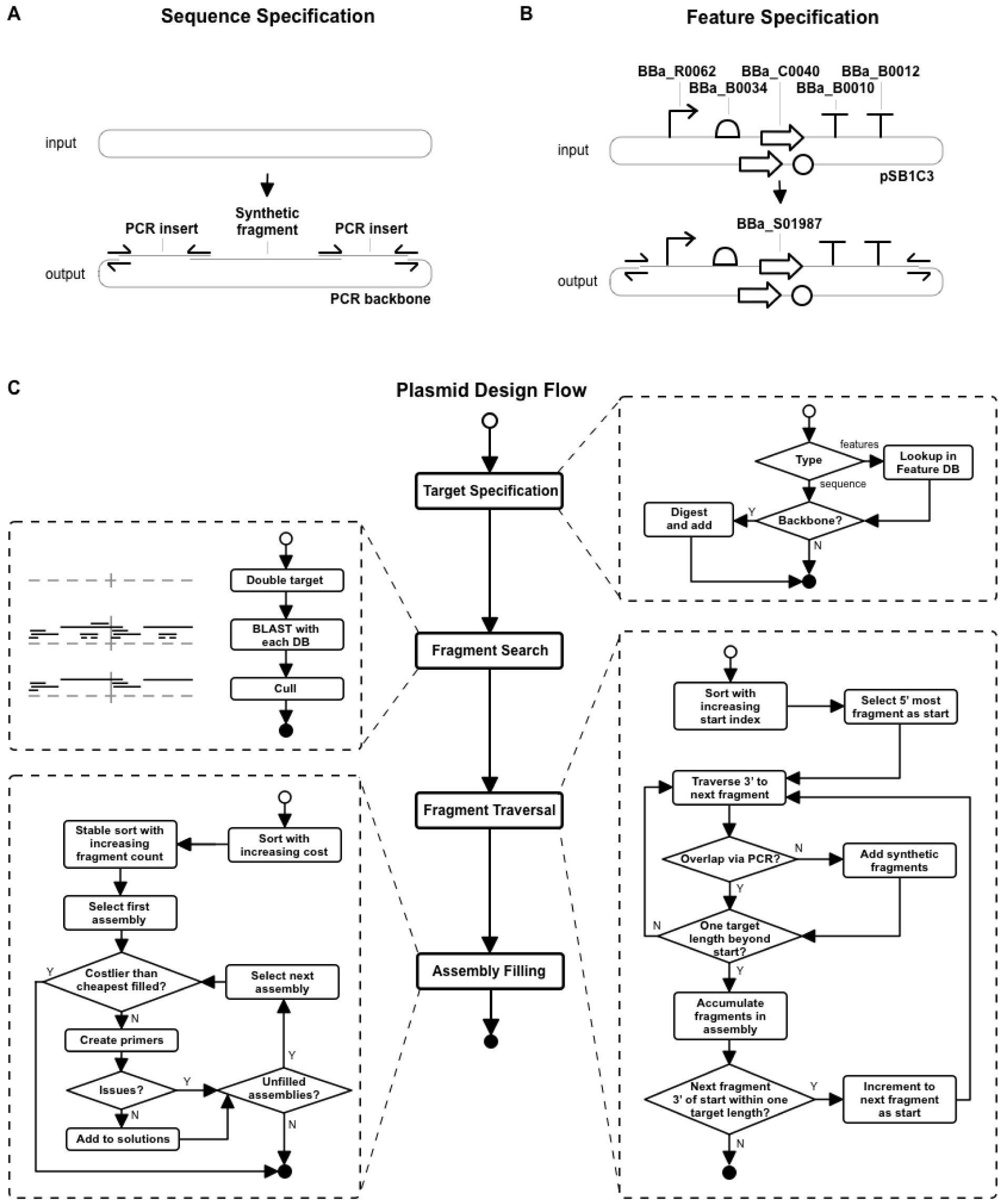
Flowchart Overview of REPP’s Plasmid Design. (A) Sequence-specified plasmid design with an example of design output with one synthetic fragment and three PCR fragments. (B) Feature-specified plasmid design with an example where iGEM parts are ordered in the orientation of the target plasmid – REPP finds a vector, BBa_S01987, that matches the user’s design specification. Users can specify their target plasmid by its sequence, composite features, or composite fragments (not shown). (C) Flowchart with the macro-microflow of REPP. Unfilled circles are the start of each process and filled circles are the end. Rounded rectangles are tasks and diamonds are conditions. The flowchart highlights the four steps of REPP’s design: parsing the target plasmid, finding fragments to cover the target, traversing the fragments to create a list of assemblies, and filling the Pareto-optimal assemblies.

### Repositories and Databases

REPP, at the time of publication, has three internal fragment databases that correspond to three sequence repositories: iGEM (11), Addgene (35), and DNASU (36). We downloaded the iGEM database from the iGEM website in XML format (37). We received both Addgene and DNASU DNA sequences, upon request, in February 2019. The Addgene database was in JSON format and the DNASU database was in Excel. We parsed all the databases to multi-FASTA format and prepared each as a BLAST database (38).

If the user specifies a target plasmid using feature names, REPP looks up each in the Feature Database – a one to one mapping between name and sequence in tab-separated values (TSV) format. REPP comes with 1096 plasmid features made public by SnapGene (24,39). On selection in a feature-based plasmid specification, each feature sequence is individually queried against the user selected databases (with a default percentage identity of 96%). Users can also choose a backbone and enzyme. Without a backbone and enzyme, REPP assumes the target is a circular plasmid, but, with them, it digests the backbone with the enzyme, selects the largest digest fragment, and anneals the target DNA sequence to the backbone’s sequence. Backbones are selected by their ID in a database and enzymes are referenced by their name in REPP’s editable Enzyme Database – another map, this one between enzymes’ names and recognition sequences (also in TSV format). Both the Feature Database and the Enzyme Database are editable from REPP’s command line interface.

### Costs

Many distinct costs go into plasmid construction and REPP factors in several of them. The most complex and variable cost is that of synthesis. REPP’s user modifiable configuration file (Supplemental Table 1), in YAML map format, supports a definition of synthesis cost curves with both variable (per basepair) and fixed costs. When REPP estimates the cost of a synthetic fragment, it looks up the first fragment integer key in the “synthetic-fragment-cost” map that exceeds the length of synthetic fragment. If the cost is fixed, it is used as is. Otherwise, the fragment’s length is multiplied by the cost (for a variable length cost). By default, the synthetic fragment cost curve and synthetic gene cost curve both correspond to IDT’s prices. Primer cost is also factored in with a default price of $0.60 per bp (IDT). Taq DNA Polymerase PCR Buffer is included at $0.27 per reaction (ThermoFisher). Gibson Assembly^®^ Master Mix is included and is estimated at $12.98 per assembly (NEB). Additionally, the cost of DNA procurement from iGEM, Addgene, and DNASU is estimated at $0, $65, and $55 per source plasmid. Addgene and DNASU’s cost per sample is straightforward, but prices are reduced for bulk orders, which REPP does not account for. Further, iGEM costs are ignored by REPP because they are bundled in iGEM teams’ registration costs or paid for once with a yearly “Labs Program” subscription (40). It assumes a $0 procurement cost for fragments in a user-defined repository.

An interesting but highly variable cost is that of “human hours”: the cost of a researcher’s time to assemble the plasmid after PCR and Gibson Assembly. There are two costs in the configuration file, one for PCR and one for Gibson Assembly, to account for these human costs – both are zero by default.

### Fragment Search

REPP finds building fragments using the target plasmid sequence doubled, end to end, as a query against each of the user selected databases. BLAST’s “blastn” task is used with reward, penalty, gapopen, and gapextend values based upon the researcher’s specified percentage identity. For example, at a percentage identity of 90% REPP uses a reward, penalty, gapopen and gapextend of 1, -2, 1, and 2, respectively. These values are documented in Supplemental File 3 and are based largely upon the BLAST user manual recommendations (41). The one exception is penalty, gapopen, and gapextend values of -6, 6, and 6 when the user specifies a percentage identity greater than 99% – we found that, to avoid imperfect hits and mismatches at the end of fragments, we needed to supply penalties that exceeded typical values for these parameters. More details are in Supplemental Table 2. After fragment matches are found and sorted along the sequence, we cull the matches to remove fragments that are fully engulfed by others. Our algorithm’s runtime is bound by BLAST and the Fragment Search step.

### Fragment Traversal

Fragments that are within one target plasmid sequence length of the start become seed points for plasmid traversal along target plasmid sequence. Candidate assemblies are accumulated during a 5’ to 3’ traversal along the target sequence from each seed, starting point. During traversal, each fragment is sequentially “annealed” into candidate assemblies with each fragment to its 3’ end. If there is a gap between fragments, the estimated number of synthetic fragments necessary to fill it is estimated. If an assembly is generated that spans the entire length of the target sequence, and has fewer fragments than the configured limit, it is stored as a valid assembly. Each fragment that starts within one target length is used, in turn, as a starting point for traversal.

### Assembly Filling

REPP’s two goals when constructing a plasmid are to minimize the design’s cost and total number of fragments. These goals usually conflict. Generally, a plasmid with large and synthetic fragments will be more expensive but require less work and preparation than a plasmid with many more fragments prepared with PCR. Because of this trade-off, REPP returns the Pareto optimal set of solutions for each target plasmid so the user can choose for themselves. During assembly filling, it sorts assemblies by their cost and then by their number of fragments. It does so to avoid non-Pareto optimal plasmid designs. If a hypothetical assembly has four fragments, but costs more than an earlier assembly that with three fragments, REPP skips the remaining assemblies to reduce execution time. The trade off in lower fragment counts for higher costs is apparent in Supplemental Fig. 2 where, when REPP designs multiple solutions, those with only two fragments are 1.7 times more expensive than the cheapest solution, on average. Other than this paragraph, when describing a plasmid’s “solution”, we are referring to the solution with the lowest cost, ignoring that REPP may have returned other more expensive solutions with fewer fragments.

When filling an assembly, REPP creates primers for the repository-derived fragments using Primer3 (42,43). By default, neighboring fragments must have at least 15 bp of homology for one another and no more than 150 bp. If adjacent fragments in an assembly will both be PCR’ed and both lack homology for one another in their source material, additional bp are included within primers’ sequences – the additional bp are equally distributed between the adjacent fragments. Conversely, when two fragments have excessive homology for one another REPP uses a subsequence one or both of the fragments to reduce the total length of overlap (Supplemental Fig 2). If there is more than the minimum amount of homology in a junction between fragments, or if one of the fragments is synthetic and the other is the product of PCR, Primer3 is given a range on the fragment’s source material so it can choose an “optimal” primer pair with more flexibility. Finally, if there is a small “gap” between two neighboring fragments that will be PCR amplified before Gibson Assembly, the template sequence will be embedded within the fragments’ primers. The default maximum length for primer-filled sequence gaps is 20 bp.

REPP checks primers for ectopic, mismatching binding sites in building fragments’ source sequences. It does so with an approach based upon Primer-BLAST (44). Primer sequences are queried with BLAST using a percentage identity of 65% and an expected value cutoff of 30,000. The annealing tm of each potential ectopic binding site found through BLAST is estimated with Primer3’s “ntthal” and “END1” mode where the first sequence is the primer and the second sequence is the reverse complement of the ectopic sequence. Assemblies are not filled if they include primers with an ectopic binding site with a primer melting temperature exceeding REPP’s “pcr-primer-max-ectopic-tm” parameter.

REPP also generates synthetic fragments during the fill step. For spans larger than the maximum synthetic fragment size limit, it splits the region into sub-fragments and creates each individually. It checks for hairpins in the junctions between the sub-fragments using “ntthal”‘s hairpin mode. If it predicts that a hairpin will form at the end of a sub-fragment and that it will exceed the “fragments-max-hairpin-tm” parameter in the configuration file, it iteratively extends the 3’ end repeatedly in an attempt to avoid it.

### Characterization with iGEM and Addgene

We tested REPP by using it to reconstruct two DNA sequence datasets: iGEM and Addgene (45,46). We specified the target plasmids’ sequences, and REPP output the repository fragments and primers to generate the plasmid’s sequence. The synthesis provider for both iGEM and Addgene datasets was IDT. For both runs the source repository was the sole source of building fragments – only iGEM was used to construct the iGEM dataset and only Addgene was used to construct the Addgene dataset. When building each dataset, we filtered out fragments from “future” years in the repository. For example, when we built a plasmid with a composite part from 2008, we excluded all iGEM parts from 2008 through present day to mirror the tool’s imagined utility to a user in 2008. Similar filtering functionality is available via an “exclude” flag in REPP’s command line interface.

The iGEM dataset contained all 24,909 parts submitted between 2005 – the year after pSB1A3 was submitted – and 2018 (11,12). We assembled each in plasmids with the backbone pSB1A3 linearized with PstI (47). The iGEM repository was the most attractive test dataset – among iGEM, Addgene, and DNASU – because all its parts can be used as insert sequences in new plasmid designs. Knowing each plasmid’s insert sequence allowed us to compare REPP’s specified assembly costs against synthesis options.

The Addgene dataset contained the 7,352 plasmids uploaded to Addgene in 2018 (Supplemental 2). Unlike the iGEM dataset, it was not explicit which parts of the plasmid were backbone and insert, so we could not compare the solutions’ costs to synthesis providers.

To test REPP’s performance with multiple synthesis providers, we created separate settings files with synthetic fragment cost curves from six synthesis providers, as of February 2019 (Supplemental Fig. 1), and rebuilt both datasets. Every REPP plasmid design solution was checked for validity by programmatically comparing the ends of adjacent fragments in the designs. If there was at least 15 and no more than 150 bp of overlap between the end of each fragment and its next a junction was considered valid. Both datasets were built with REPP on a Google Compute instance with 16 virtual cores and Ubuntu 16.

## RESULTS

After building iGEM plasmids with DNA fragments from the iGEM repository and/or synthetic fragments, we estimated the total assembly costs to build the iGEM dataset via REPP, synthetic fragments, and synthetic genes to be $3,594,327, $5,411,060 and $12,854,997. In other words, if the entire dataset was assembled, REPP would save 34% versus synthetic fragments alone and 72% versus synthetic genes. Approximately 35% of plasmids had at least 30% cost savings compared to synthetic fragments alone (Fig 2C).

The length of the inserted iGEM part greatly impacted cost savings (Fig 2F). While REPP reverted to synthesis for almost all small plasmids less than 500bp, its savings versus synthetic fragments alone improved to an average of at least 37% for all plasmids with iGEM parts longer than 3,000bp. This makes intuitive sense: there were more opportunities for sequence re-use from existing parts in the iGEM repository.

After building the Addgene dataset, the median cost for Addgene plasmid designs for Addgene was $374 versus $155 for the iGEM dataset. The difference can be explained by the standard linearized backbone (pSB1A3) used for the iGEM datasets – without it in the Addgene dataset, REPP had a greater amount of variable sequence to cover in each plasmid.

Choice in synthesis provider affected REPP’s solution costs. For the iGEM dataset, the plasmid cost quartiles were $65, $85, and $125 versus $102, $162, and $233 from Twist and IDT, respectively (Fig 3A). It should be noted that REPP does not, yet, encode or check for synthesis complexities from different synthesis providers. The probability and time of procurement may vary between synthesis providers but was not factored into costs or plasmid designs. There are other tools like BOOST that support sequence redesigns to improve synthesis favorability (18,30).

Percentage identity also affected price (Fig 3C, Fig 3D). The mean price declined from $222 to $206 for iGEM and from $695 to $540 for Addgene when the percentage identity of the resulting plasmid was lowered from 100% to 98%. A lower identity, with its concomitant loosening of BLAST parameters, led to REPP finding a greater number of potential building fragments and cheaper overall solutions. Because of the drop-off in cost at 98% identity, we used it as the default percentage identity; though it is adjustable through REPP’s CLI.

The median build time for the iGEM dataset was 0.46 ± 1.4 seconds while the median build time for the Addgene dataset was 15 ± 8.9 seconds; BLAST was the bottleneck in execution times for both datasets.

## DISCUSSION

We have demonstrated that plasmid design benefits from pairing with DNA repositories. We have made and characterized an open-source application, REPP, that finds building fragments in public and user-specific DNA repositories. It prepares repository fragments for Gibson Assembly via PCR with Primer3 primers and synthesis is considered in all assemblies, so the least expensive possible assembly is generated given a set of design rules. In particular, REPP performs best on larger plasmids with inserts greater than 1,000bp where the probability of sequence re-use is greatest.

REPP’s scope could be expanded in the future. First, the approach described here is specific to Gibson Assembly but could be adapted for other cloning methods. For example, sets of building fragments that have the requisite fusion sites for a complete Golden Gate assembly could be used without modification; otherwise, fragments could be prepared for Golden Gate via primers with embedded BsaI or BbsI sites (31). Second, genomes and additional sequence repositories could augment REPP’s list of build fragment sources via a remote database management system like SynBioHub (32) – users could specify the genome or public repository they would like to include as a build source and those sources would be downloaded on command. Third, REPP could be expanded to include time-to-build optimization via integration with synthesis providers’ web-APIs for information on synthetic DNA procurement times. Finally, REPP’s bio-design capabilities could be augmented via pairing with higher-level tools like Eugene (33), Cello (2), and Double Dutch (34).

We believe that the sequence or feature based plasmid design approach described here is more intuitive that existing state-of-the-art fragment-based design tools and that it will enable complex and multi-source plasmid designs that would be infeasible otherwise.

## ACKNOWLEDGEMENTS

We thank Kevin LeShane (Lattice Automation) for comments on the manuscript.

